# DEAD-box protein family member DDX28 is a negative regulator of HIF-2α and eIF4E2-directed hypoxic translation

**DOI:** 10.1101/632331

**Authors:** Sonia L. Evagelou, Olivia Bebenek, Erin J. Specker, James Uniacke

## Abstract

Hypoxia occurs when there is a deficiency in oxygen delivery to tissues and is connected to physiological and pathophysiological processes such as embryonic development, wound healing, heart disease and cancer. The master regulators of oxygen homeostasis in mammalian cells are the heterodimeric hypoxia-inducible transcription factors HIF-1 and HIF-2. The oxygen-labile HIF-2α subunit has not only been implicated in transcription, but also as a regulator of eIF4E2-directed hypoxic translation. Here, we have identified the DEAD-box protein family member DDX28 as a novel interactor and negative regulator of HIF-2α that suppresses its ability to activate eIF4E2-directed translation. We demonstrate that stable silencing of DDX28 via shRNA in hypoxic human U87MG glioblastoma cells caused an increase, relative to control, to: HIF-2α protein levels, the ability of eIF4E2 to bind the m^7^GTP cap structure, and the translation of select eIF4E2 target mRNAs. DDX28 depletion elevated both nuclear and cytoplasmic HIF-2α, but HIF-2α transcriptional activity did not increase possibly due to its already high nuclear abundance in hypoxic control cells. Depletion of DDX28 conferred a proliferative advantage to hypoxic, but not normoxic cells, which is likely a consequence of the translational upregulation of a subset of hypoxia-response mRNAs. DDX28 protein levels are reduced in several cancers, including glioma, relative to normal tissue. Therefore, we uncover a regulatory mechanism for this potential tumor suppressor in the repression of HIF-2α- and eIF4E2-mediated translation activation of oncogenic mRNAs.

## INTRODUCTION

The procurement for oxygen is a fundamental aspect of survival for aerobic organisms in all domains of life. Mammals have evolved complex circulatory, respiratory, and neuroendocrine systems to satisfy the need for molecular oxygen as the primary electron acceptor in oxidative phosphorylation, which supplies energy in the form of ATP [1]. The ability of a cell to acclimate to low oxygen (hypoxia or ≤ 1% O_2_), which usually arises due to an imbalance in supply and demand, is essential from the earliest stages of life [2, 3]. Hypoxia plays a role in several physiological and pathophysiological conditions such as embryonic development, muscle exercise, wound healing, cancer, heart disease, and stroke [4]. Prior to the establishment of uteroplacental circulation, embryonic cells receive only as much as 2% O_2_, and following oxygenation by maternal blood, the embryo still contains discrete regions of hypoxia [2, 5]. This form of physiological hypoxia helps govern the process of development through cell fate determination, angiogenesis, placentation, cardiogenesis, bone formation, and adipogenesis [3, 6–10]. Hypoxia is also a feature of the tumor microenvironment, and plays a key role in several cancer hallmarks toward tumor progression [11, 12]. The major cellular response to hypoxia is mediated by hypoxia-inducible transcription factors (HIF-1 and HIF-2). The HIFs are heterodimeric, composed of an oxygen-labile α-subunit, HIF-1α or HIF-2α, and a constitutively expressed HIF-1β subunit [11]. HIF-1 and HIF-2 activate the transcription of hundreds of genes (some shared and some unique), including those involved in metabolism and erythropoiesis, in order to simultaneously reduce the activity of energy-expensive processes and promote the increased uptake of nutrients and oxygen [13]. Further investigation into the unique roles of HIF-1α and HIF-2α, and how they might be differentially regulated, will reveal novel insights into hypoxic gene expression.

Unlike HIF-1α that is more involved in acute hypoxia (< 24 h), HIF-2α is linked to chronic hypoxia [14]. Further, HIF-2α accumulates in both the cytoplasm and the nucleus [15]. Indeed, HIF-2α, but not HIF-1α, participates in the recruitment of select hypoxia-response mRNAs to the translation apparatus [16], not only in hypoxia but in the low-range of physiological oxygen (< 8% O_2_) [17]. Hypoxia is a potent inhibitor of mammalian target of rapamycin complex 1, suppressing canonical cap-dependent translation by sequestering the cap-binding protein eukaryotic initiation factor 4E (eIF4E) [18–21]. Alternative modes of translation initiation are utilized during hypoxia including cap-independent mechanisms as well as non-canonical cap-dependent translation mediated by the eIF4E2 cap-binding protein. HIF-2α and RBM4 recognize select mRNAs by binding to RNA Hypoxia Response Elements (rHREs) in their 3’ UTRs and initiating their translation via the 5’ cap through eIF4E2, eIF4G3 and eIF4A [16, 22]. HIF-2α and eIF4E2 are essential for embryonic development [3, 23], and important contributors to tumor progression [24, 25], both hypoxia-driven processes. The protein levels of eIF4E2 do not change between normoxia and hypoxia [16, 17]. HIF-2α, on the other hand, is constantly degraded in normoxia via a family of prolyl hydroxylases that are inhibited by hypoxia [11]. In hypoxia, it is mostly unclear how the activities of eIF4E2 or HIF-2α are regulated with respect to translation initiation. A greater understanding of these mechanisms will shed light on how gene expression is coordinated in physiological and pathophysiological processes that are linked to hypoxia.

Here we demonstrate that the DEAD box protein DDX28 is a negative regulator of HIF-2α protein levels in a human glioblastoma cell line. DDX28 has been detected in mitochondria, cytoplasm and nuclei [26]. However, known functions of DDX28 are limited to its RNAi-mediated silencing disrupting mitoribosome assembly [27, 28], which is also an outcome of hypoxia [29]. While it is known that hypoxia increases HIF-2α levels, our data show that DDX28 protein levels concurrently decrease. We also show that DDX28 interacts with HIF-2α, but not HIF-1α or the m^7^GTP cap structure. We demonstrate a link between these two observations by depleting DDX28 levels in hypoxia whereby an even greater increase to HIF-2α levels was observed along with its effect on eIF4E2-dependent translation activation. Hypoxic depletion of DDX28 caused eIF4E2 and its mRNA targets to associate more with the m^7^GTP cap structure and polysomes, respectively. Furthermore, hypoxic depletion of DDX28 caused a significant increase to cell proliferation only in hypoxia. While HIF-2α increased in both the nucleus and cytoplasm upon DDX28 depletion, there was no increase to its transcriptional activity. We propose a model where DDX28 reduction in hypoxia plays a role in HIF-2α stabilization, but some DDX28 is still useful to restrain the HIF-2α/eIF4E2 translational axis, which can be oncogenic [24, 25].

## RESULTS

### DDX28 interacts with HIF-2α, but not HIF-1α or eIF4E2

U87MG human glioblastoma cells were used in this study because they have previously been characterized as models for hypoxia research and the interaction between HIF-2α and eIF4E2 in non-canonical cap-dependent translation [16, 17, 22, 24, 25]. When U87MG were exposed to hypoxia (1% O_2_) for 24 h, we not only observed an increase in HIF-2α, but a concurrent decrease in DDX28 levels (Fig. 1A). Furthermore, when DDX28 was stably depleted via two independent shRNA sequences to produce two unique cell lines, the hypoxia-dependent increase in HIF-2α was further increased relative to controls expressing a non-targeting shRNA (Fig. 1B). We next investigated whether DDX28 was directly involved in HIF-2α regulation by performing a co-immunoprecipitation. Exogenous tagged proteins were used due to the lack of specificity of the antibodies for the endogenous protein or its low abundance. In hypoxic U87MG cells, exogenous GFP-HIF-2α, but not GFP alone, co-immunoprecipitated with DDX28 (Fig. 1C). Since HIF-2α is a known interactor of eIF4E2 at the 5’ mRNA cap in hypoxia [16], we tested whether DDX28 was part of this complex. However, exogenous eIF4E2 co-immunoprecipitated with HIF-2α, but not DDX28 (Fig. 1D). Further, to demonstrate specificity for the HIF-2α homolog, HIF-1α did not co-immunoprecipitate with DDX28 (Fig. 1E). These data suggest that DDX28 interacts with a distinct pool of HIF-2α that is not associated with eIF4E2.

**Figure 1.**
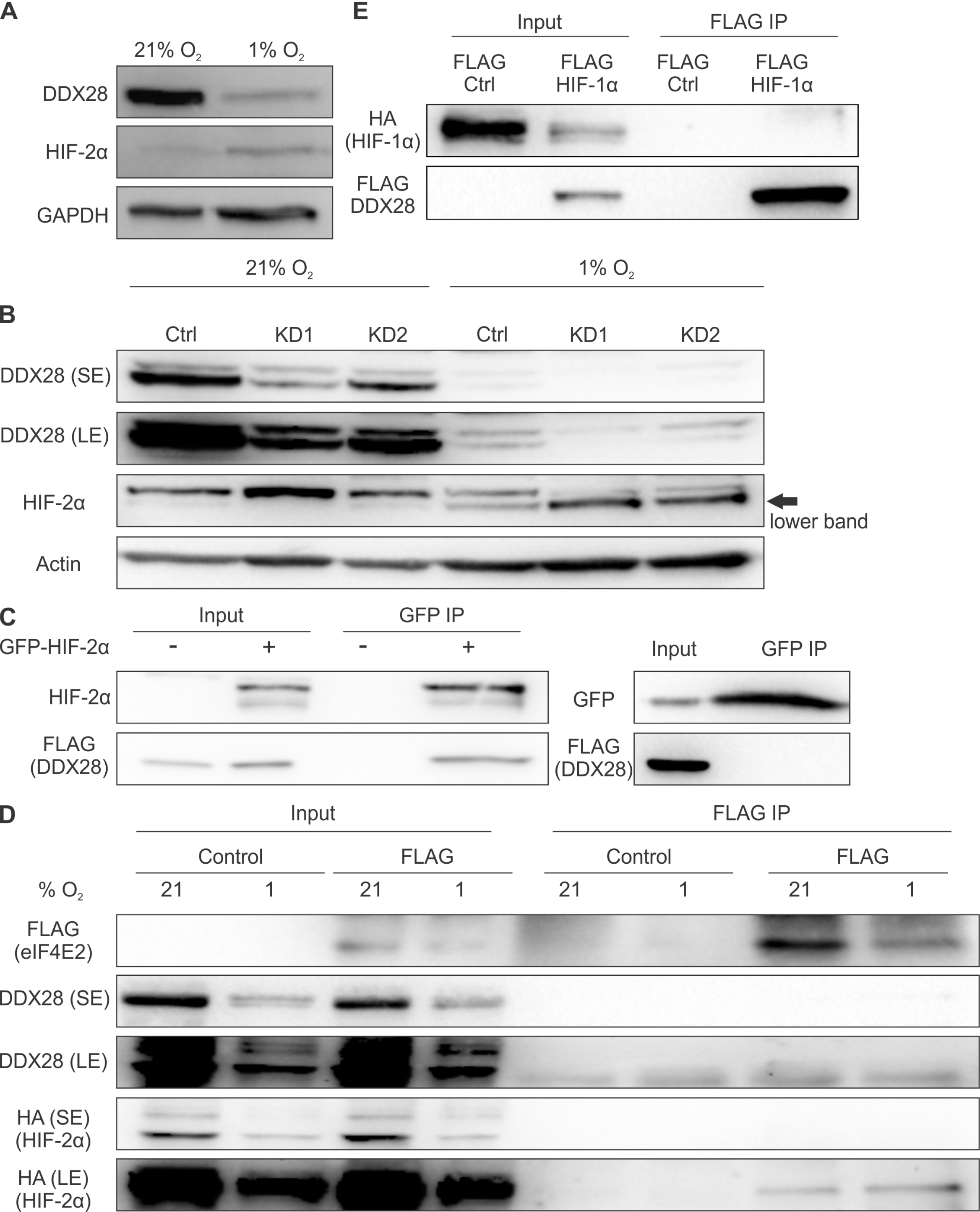
DDX28 interacts with HIF-2α, but not HIF-1α or eIF4E2. (A) Western blot of total HIF-2α and DDX28 protein levels in normoxia (21% O_2_) and hypoxia (1% O_2_). GAPDH used as a loading control. (B) Western blot of DDX28 and HIF-2α (arrow, hypoxia-inducible lower band) normoxic and hypoxic protein levels in control (Ctrl) cells stably expressing a non-targeting shRNA or in cells stably expressing one of two shRNAs targeting DDX28 mRNA: Knockdown (KD) 1 and KD2. Actin used as a loading control. (C) GFP co-immunoprecipitation in hypoxic cells stably expressing FLAG-DDX28 and transfected with recombinant GFP-HIF-2α. Cells transfected with no DNA (−) or GFP alone were used as controls. (D) FLAG co-immunoprecipitation of recombinant FLAG-eIF4E2 from hypoxic cells co-transfected with HA-HIF-2α. Cells transfected with empty FLAG vector used as control. (E) FLAG co-immunoprecipitation in hypoxic cells stably expressing FLAG-DDX28 and transfected with recombinant HA-HIF-1α. Cells transfected with empty FLAG vector used as control. SE, short exposure; LE, long exposure. 25 μg of whole cell lysate was used as input. Experiments performed in U87MG glioblastoma.

### Depletion of DDX28 enhances the association of eIF4E2 with m^7^GTP and polyribosomes

We performed m^7^GTP cap-binding assays to test whether the increased HIF-2α levels in DDX28 depleted cells had an effect on the translation initiation potential of eIF4E2. In hypoxia, eIF4E2 bound more to m^7^GTP by 1.6 ± 0.1 and 1.5 ± 0.2 fold in two DDX28 depleted cell lines relative to control (Fig. 2A-B). Surprisingly, depletion of DDX28 in normoxia had an even greater effect on eIF4E2 binding to m^7^GTP with 2.9 ± 0.4 and 2.5 ± 0.1 fold increases relative to control (Fig. 2C-D). These data are also in support of Fig. 1D that DDX28 does not bind the m^7^GTP cap.

**Figure 2.**
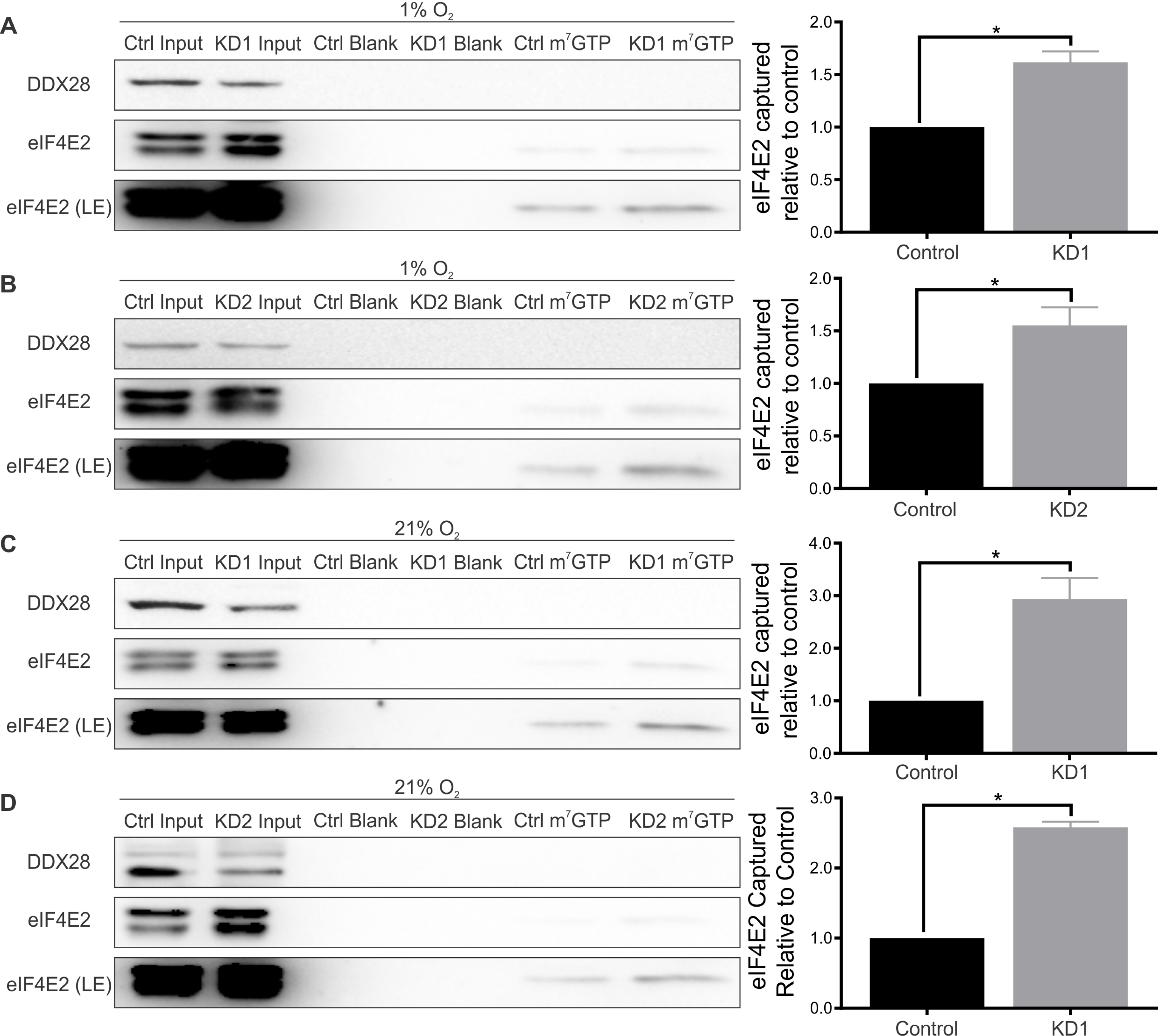
Knockdown of DDX28 increases the cap-binding affinity of eIF4E2. Western blot and quantification of eIF4E2 capture with m^7^GTP-bound agarose beads in cells stably expressing one of two distinct shRNA sequences targeting DDX28 in 1% O_2_ hypoxia (A-B) and 21% O_2_ normoxia (C-D). 35 μg of whole cell lysate was used as the input. Ctrl, control cells stably expressing non-targeting shRNA; KD1 and KD2, knockdown cells stably expressing one of two distinct shRNA sequences targeting DDX28; LE, long exposure. Data (n ≥ 3), mean ± s.e.m normalized to input. * represents p < 0.05 using one sample t-test against hypothetical mean (μ = 1). Experiments performed in U87MG glioblastoma.

m^7^GTP association of a cap-binding protein like eIF4E2 does not necessarily imply an increase in translation initiation [30]. Therefore we performed polysome fractionation to test whether eIF4E2 had a greater association with polysomes isolated from DDX28 depleted cells relative to controls in normoxia and hypoxia. Using densitometry to quantify total eIF4E2 associated with monosomes (low translation; fractions 1-3) and polysomes (medium to high translation; fractions 4-9), we show that 5 ± 1.2% of total eIF4E2 in normoxic control cells was associated with polysomes (Fig. 3A). In DDX28-depleted cells, the proportion of polysome-associated eIF4E2 increased to 23 ± 3.2% (Fig. 3B). Hypoxic control cells displayed 22 ± 5.1% eIF4E2 associated with polysomes (Fig. 3C), an increase relative to normoxic control cells (5 ± 1.2%; Fig. 3A). Hypoxic DDX28-depleted cells had an even greater proportion of eIF4E2 associated with polysomes (41 ± 4.1%) relative to hypoxic control cells (Fig. 3D). The polysome association of the canonical cap-binding protein eIF4E did not change in any condition (Fig. 3E). Our data show that hypoxia, or knockdown of DDX28 in normoxia, increased the proportion of polysome-associated eIF4E2 relative to normoxic control cells (Fig. 3F). However, DDX28 depletion and hypoxia together significantly increased the polysome association of eIF4E2 relative to normoxic control cells (Fig. 3F).

**Figure 3.**
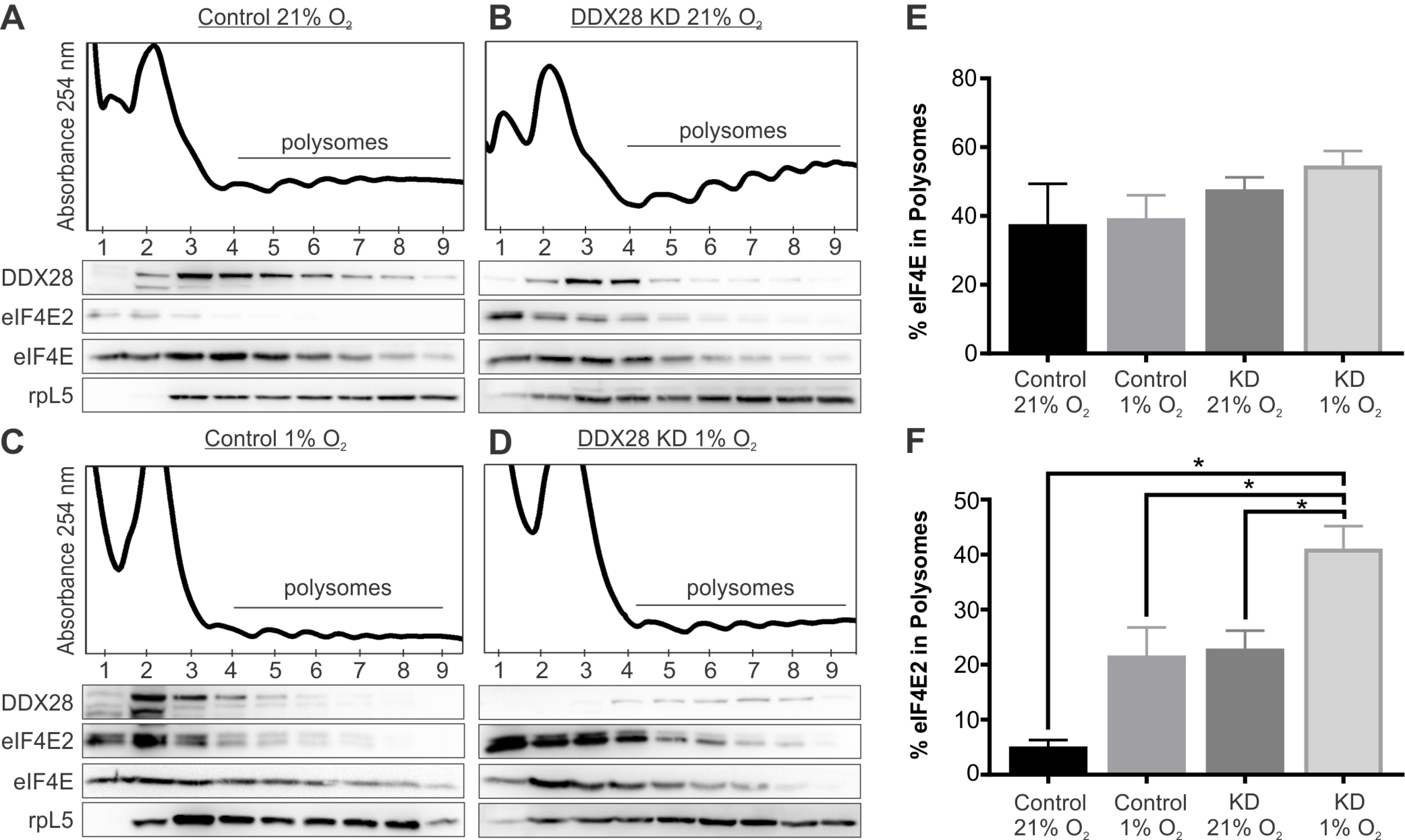
Knockdown of DDX28 increases polysome-associated eIF4E2 only in hypoxia. Polysomal distribution of DDX28, eIF4E2 and eIF4E protein measured by western blot in control cells stably expressing non-targeting shRNA in 21% O_2_ normoxia (A) and 1% O_2_ hypoxia (C) and in Knockdown (KD) cells stably expressing an shRNA targeting DDX28 in normoxia (B) and hypoxia (D). Ribosomal protein L5 (rpL5) used as a marker of protein integrity in each fraction. The eIF4E (E) or eIF4E2 (F) protein associated with polysomes (fractions 4-9) as a percentage of total protein (fractions 1-9) was quantified by densitometry. Data (n = 3), mean ± s.e.m. * represents p < 0.05 using one-way ANOVA and Tukey’s HSD post-hoc test. Experiments performed in U87MG glioblastoma.

### Depletion of DDX28 increases the translation of eIF4E2 target transcripts in hypoxia

We performed qRT-PCR on monosome and polysome fractions in normoxia and hypoxia to measure the DDX28-dependent association of eIF4E2 and eIF4E target transcripts. We chose Epidermal Growth Factor Receptor (*EGFR*) and Insulin Like Growth Factor 1 Receptor (*IGF1R*) mRNAs, previously characterized as eIF4E2-dependent transcripts due to the presence of an rHRE in their 3’ UTR [16, 17], and Eukaryotic Translation Elongation Factor 2 (*EEF2*) and Heat Shock Protein 90 Alpha Family Class B Member 1 (*HSP90ab1*) mRNAs previously characterized as eIF4E-dependent transcripts due to the presence of a 5′ terminal oligopyrimidine motif [17, 31]. We observed a significant increase in the association of *EGFR* mRNA with polysomes relative to monosomes from 4.5 ± 0.9-fold in controls to 11.3 ± 1.6-fold in DDX28 depleted cells in hypoxia (Fig. 4A). Similarly, the polysome association of *IGF1R* mRNA significantly increased from 3.2 ± 0.3 in controls to 5.4 ± 0.7 in DDX28-depleted cells in hypoxia (Fig. 4A). In normoxia, there was no difference in the polysome-association of *EGFR* mRNA relative to monosomes between controls (3.5 ± 0.5-fold) and DDX28 depleted cells (3.8 ± 0.5-fold) (Fig. 4B). Similarly, there was no statistical difference in the polysome-association of *IGF1R* mRNA relative to monosomes between controls (5.6 ± 0.8-fold) and DDX28 depleted cells (3.5 ± 0.5-fold) (Fig. 4B). Moreover, EGFR protein levels (Fig. 4C), but not total *EGFR* mRNA levels (Fig. 4D), increased in hypoxic DDX28-depleted cells relative to control. Neither of the eIF4E-dependent transcripts displayed significant changes in polysome-association in normoxia or hypoxia between DDX28-depleted and controls. The one exception was *EEF2* mRNA, which displayed a small significant increase in normoxic polysome-association relative to monosomes in DDX28-depleted cells (1.52 ± 0.04) compared to controls (1.05 ± 0.06) (Fig. 4E). These data suggest that DDX28 depletion significantly increases the translation of eIF4E2-dependent, but not eIF4E-dependent, transcripts in hypoxia.

**Figure 4.**
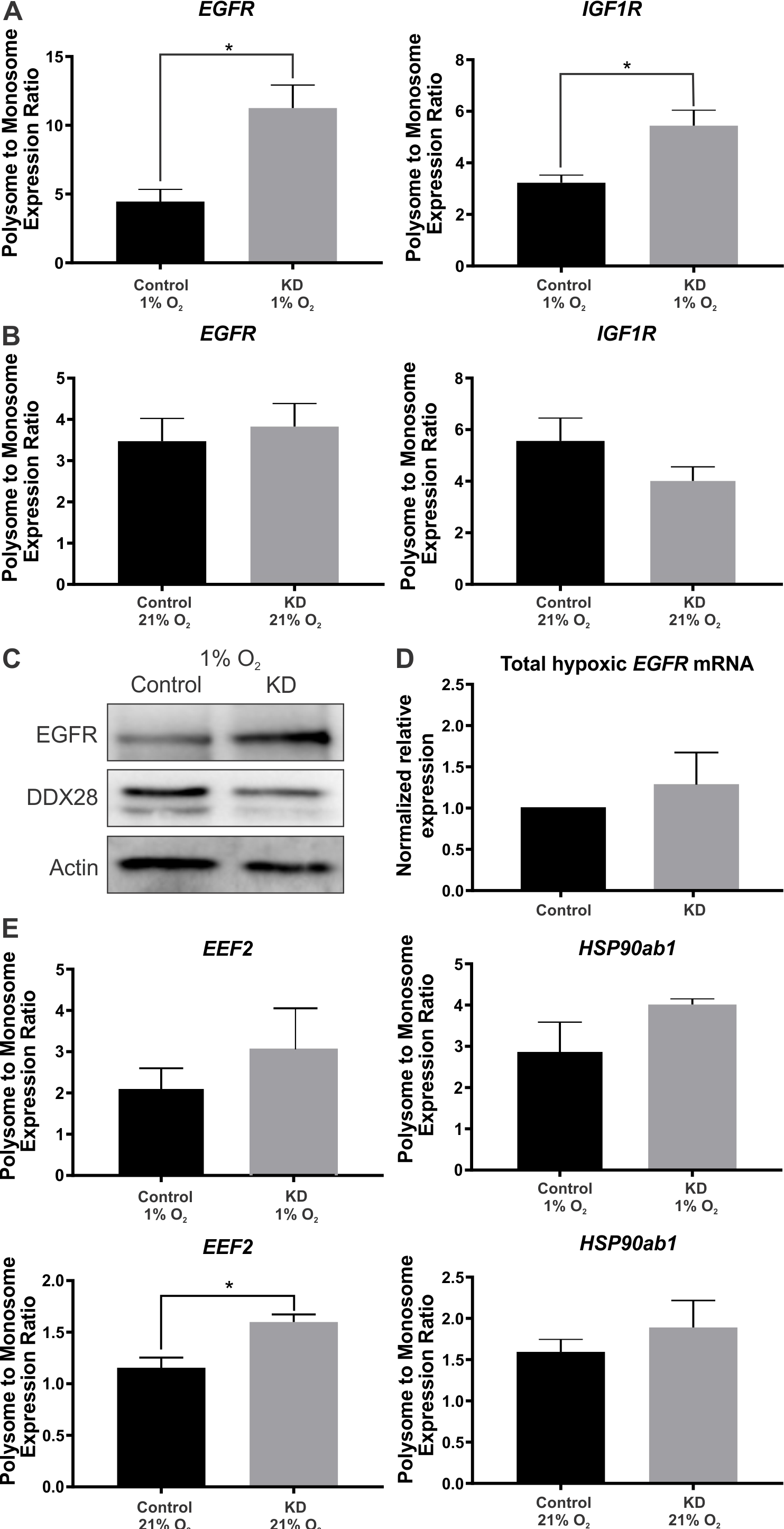
Depletion of DDX28 increases the polysome association of eIF4E2-dependent transcripts in hypoxia. The association of eIF4E2-dependent transcripts Epidermal Growth Factor Receptor (EGFR) and Insulin Like Growth Factor 1 Receptor (IGF1R) with polysomes (fractions 4-9 from Fig. 3) was measured relative to monosomes (fractions 1-3) by qRT-PCR in control cells expressing a non-targeting shRNA or Knockdown (KD) cells stably expressing an shRNA targeting DDX28 in 1% O_2_ hypoxia (A) and 21% O_2_ normoxia (B). Western blot of total EGFR, DDX28, and Actin protein levels (C) or qRT-PCR of total EGFR mRNA levels normalized to endogenous control genes RPLP0 and RPL13A (D) in hypoxic control cells or DDX28 KD cells. (E) The association of eIF4E-dependent transcripts Eukaryotic Translation Elongation Factor 2 (EEF2) and Heat Shock Protein 90 Alpha Family Class B Member 1 (HSP90ab1) with polysomes was measured relative to monosomes by qRT-PCR in control cells or DDX28 KD cells in hypoxia and normoxia. Data (n ≥ 3), mean ± s.e.m. * represents p < 0.05 using unpaired two-sample t-test except in (D) a one-sample t-test against a hypothetical mean (μ = 1).

### Depletion of DDX28 in hypoxia increases cytoplasmic and nuclear HIF-2α levels, but not its nuclear activity

Hypoxic cells were fractionated into cytoplasm and nuclei to measure the effects of DDX28 depletion on HIF-2α levels in both compartments. In accordance with the abovementioned observations that DDX28 depletion increases total HIF-2α protein levels and its cytoplasmic activity (translation), we show that DDX28 depletion increased cytoplasmic HIF-2α levels compared to control (Fig. 5A). However, we also observed an increase in the nuclear levels of HIF-2α. Therefore, we investigated whether DDX28 depletion in hypoxia also increased the nuclear activity of HIF-2α (transcription) by measuring the mRNA abundance from its gene targets relative to control. We chose six genes that contain Hypoxia Response Elements in their promoters that are more dependent on HIF-2α than HIF-1α [32–35]. None of the six genes displayed a significant increase in mRNA abundance in DDX28 depleted cells relative to control (Fig. 5B). In fact, two genes displayed significant decreases in mRNA abundance, albeit by two-fold at most. These data suggest that while both nuclear and cytoplasmic HIF-2α levels increase in response to DDX28 depletion, the effect on HIF-2α transcriptional activity is minimal.

**Figure 5.**
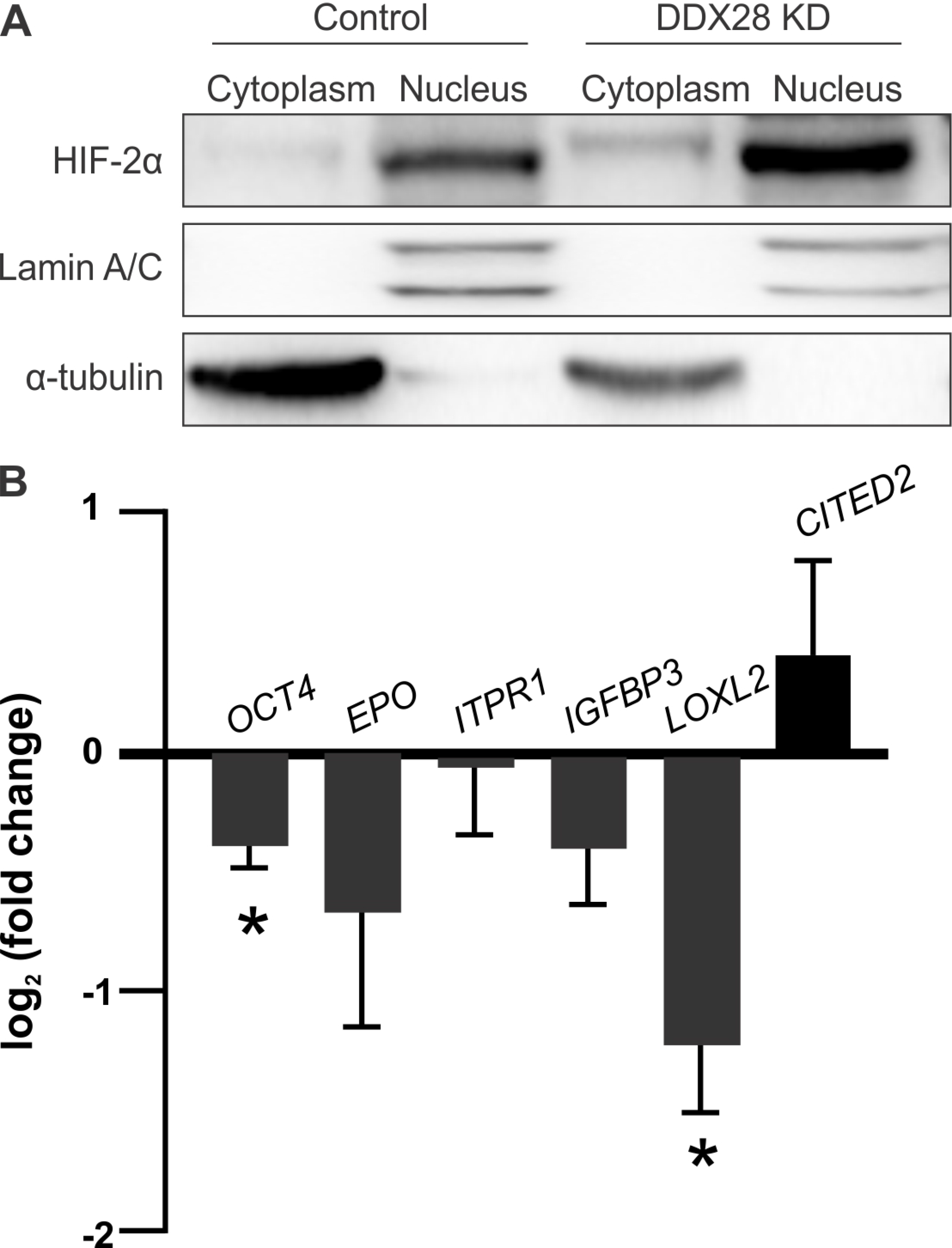
Depletion of DDX28 in hypoxia increases cytoplasmic and nuclear HIF-2α levels, but not its nuclear activity. (A) Western blot of HIF-2α protein levels in cytoplasmic and nuclear fractions of control cells expressing a non-targeting shRNA or Knockdown (KD) cells stably expressing an shRNA targeting DDX28 in 1% O_2_ hypoxia. Lamin a/c used as nuclear marker and α-tubulin as cytoplasmic marker. (B) The mRNA abundance of HIF-2α gene targets in hypoxia measured via qRT-PCR. Data (n ≥ 3), mean ± s.e.m. represented as log_2_ (fold change) in DDX28 KD cells relative to control cells and normalized to endogenous control genes RPLP0 and RPL13A. * represents p < 0.05 using a one-sample t-test against hypothetical mean (μ = 0). CITED2, Cbp/P300-Interacting Transactivator 2; EPO, Erythropoietin; IGFBP3, Insulin Like Growth Factor Binding Protein 3; ITPR1, inositol 1,4,5-trisphosphate receptor type 1; LOXL2, Lysyl Oxidase Like 2; OCT4, octamer-binding transcription factor 4. Experiments performed in U87MG glioblastoma.

### DDX28 depletion causes an increase in cell viability and proliferation in hypoxia but not normoxia

We next investigated whether the increase in eIF4E2-directed translation in DDX28 depleted cells provided a benefit to cells by measuring their viability and proliferation in normoxia and hypoxia. To assess this, we monitored the number of viable DDX28 depleted and control cells over 72 h at 24 h intervals using crystal violet staining. We observed no significant differences in viability at 24, 48, or 72 h between normoxic DDX28 depleted and control cells (Fig. 6A). However, both hypoxic DDX28 depleted cell lines had significantly increased viability compared to control at each time interval (Fig. 6B). To measure proliferation, we monitored the incorporation of bromodeoxyuridine (BrdU) into the DNA of actively dividing cells via immunofluorescence. In normoxia, one DDX28 depleted cell line displayed a significant increase in BrdU incorporation relative to control (Fig. 6C). However, following 24 h of hypoxia, both DDX28 depleted cell lines displayed significant increases in BrdU incorporation relative to control (Fig. 6D). To test whether overexpressing exogenous DDX28 would have the opposite effect (decreased viability, proliferation, and HIF-2α levels) in hypoxia, we generated two stable clonal U87MG cell lines expressing FLAG-DDX28 and a control expressing FLAG alone. We did not observe any significant differences in viability, proliferation, or HIF-2α levels between the overexpressing cell lines relative to the control (Fig. S1A-E). The one exception was a decrease in viability after 24 h of hypoxia in one of the overexpressing cell lines relative to control, but this difference ceased at 48 h and 72 h. Since hypoxia appears to reduce the levels of exogenous DDX28 (Fig. S1E), it is possible that the overexpressing cell lines do not overexpress DDX28 enough in hypoxia to suppress HIF-2α or that DDX28 is already suprastoichiometric to HIF-2α. Our data suggest that depletion of DDX28 provides an increase to cell viability and proliferation in hypoxia likely through an increase in translation of select mRNAs and perhaps other unidentified pathways.

**Figure 6.**
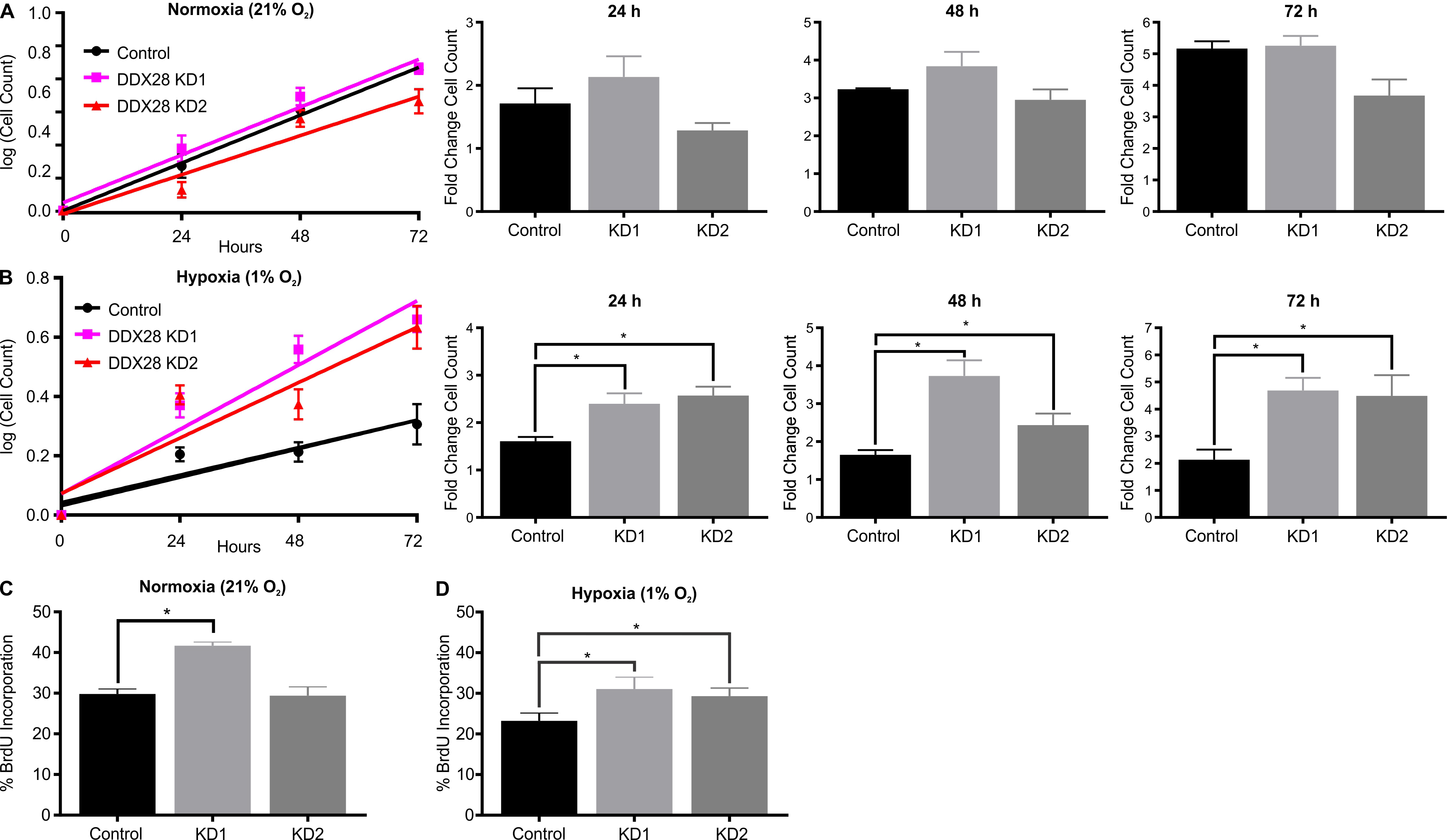
DDX28 depletion causes an increase in cell viability and proliferation only in hypoxia. Viable cell counts were measured with crystal violet staining after 24 h, 48 h, and 72 h in 21% O_2_ normoxia (A) and 1% O_2_ hypoxia (B) for control cells expressing a non-targeting shRNA or Knockdown cells stably expressing one of two shRNAs targeting DDX28 (KD1 and KD2). All absolute cell count values were normalized to the number of cells present on day 0 for each individual independent experiment, representing the fold change in the number of cells at each time point relative to day 0. Proliferation was measured as % BrdU-positive control and DDX28 KD cells via immunofluorescence after 24 h in normoxia (C) or hypoxia (D). Data (n ≥ 3), mean ± s.e.m. * represents p < 0.05 using an unpaired two-sample t-test. Experiments performed in U87MG glioblastoma.

## DISCUSSION

HIF-2α and eIF4E2 contribute to the ability of a cell to adapt to hypoxic conditions through selective gene expression by transcriptional and translation regulation. We have previously shown that eIF4E2 knockdown represses hypoxic cell proliferation and survival, migration and invasion, and tumor growth [24, 25]. Furthermore, eIF4E2-directed translation is active in the low range of physiological oxygen where HIF-2α, but not yet HIF-1α, is stabilized (3-8% O_2_) [17]. Total levels of eIF4E2 protein are minimally altered upon hypoxic exposure [16, 17, 25, 36], so how is it regulated? Post-translational modifications of eIF4E2 have been identified such as ISGylation [37], as well as protein interactors such as eIF4G3 and eIF4A [16, 22]. However, the stabilization of HIF-2α is essential for eIF4E2 hypoxic activity [16] and is likely a more upstream regulatory step for eIF4E2-directed translation to be functional.

We have uncovered a new mode of HIF-2α regulation that affects the activity of hypoxic translation via eIF4E2. HIF-2α co-immunoprecipitated with DDX28 and eIF4E2 (Fig. 4C), but eIF4E2 did not co-immunoprecipitate with DDX28 (Fig. 4D). This suggests that DDX28 interacts with a different pool of HIF-2α than the one that interacts with eIF4E2. In agreement, DDX28 did not interact with the m^7^GTP cap structure (Fig. 2). However, the depletion of DDX28 did significantly increase the ability of eIF4E2 to associate with m^7^GTP, suggesting that its effect on HIF-2α influences eIF4E2 activity. Indeed, eIF4E2 did associate more with polysomes upon DDX28 depletion (Fig. 3F). Mechanisms were initially proposed where HIF-2α acted at the 3’ UTR rHRE of select mRNAs along with RBM4 to mediate joining of the 5’ end and eIF4E2 [16, 22, 38]. It is important to note that cap-binding assays are performed with m^7^GTP bound to agarose beads and not to an mRNA [39]. Therefore, these data suggest that HIF-2α could act directly on eIF4E2 to regulate its cap-binding potential (Fig. 7).

**Figure 7.**
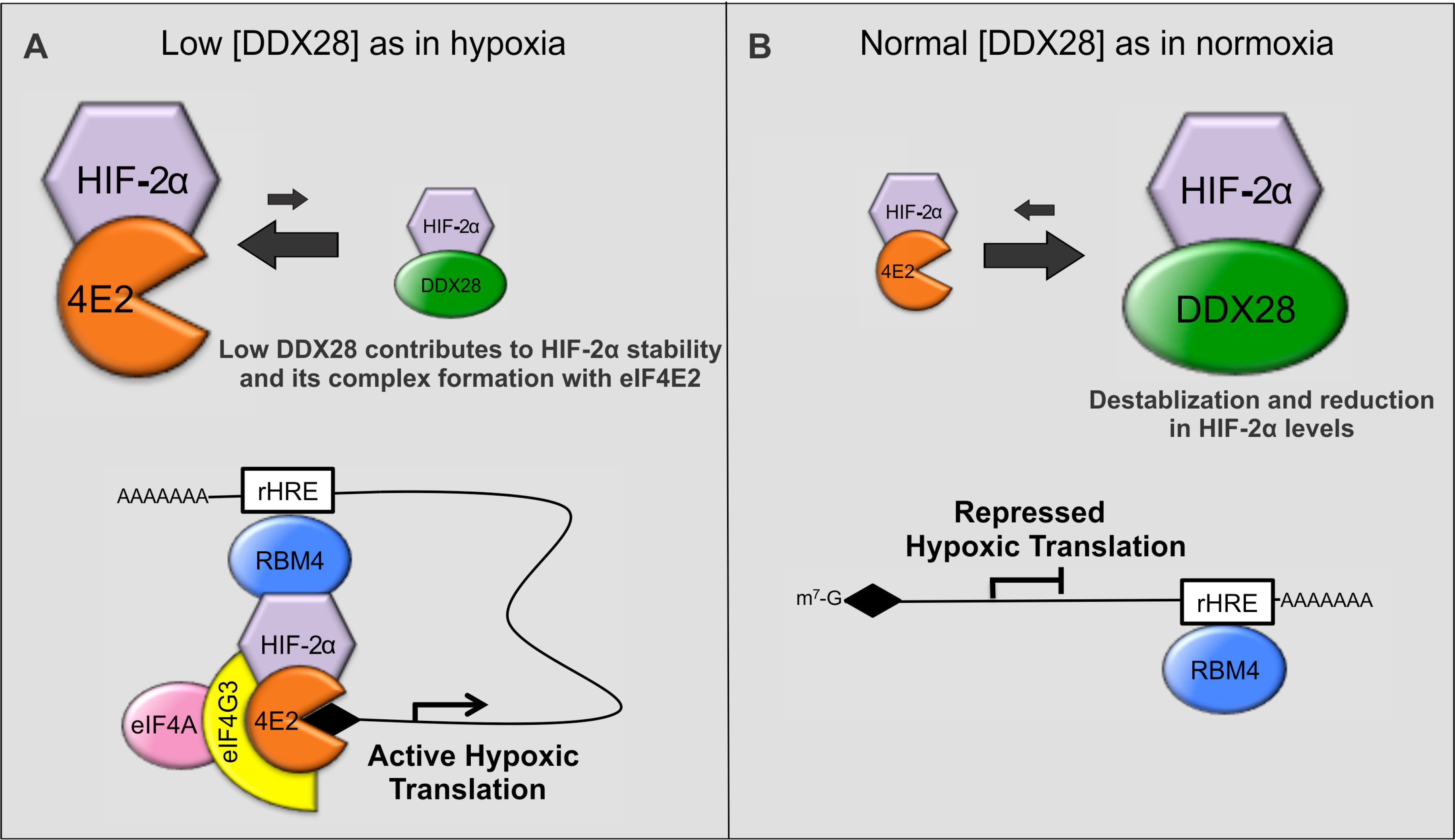
Model of HIF-2α regulation via DDX28. (A) In situations of low DDX28 such as 1% O_2_ hypoxia or shRNA depletion of DDX28 in normoxia, HIF-2α becomes more stabilized and shifts the balance toward complex formation with eIF4E2 and a higher affinity for the m^7^GTP 5’ cap structure. This, along with the binding of known interactors eIF4G3, eIF4A and RBM4, induces the translation of transcripts harboring RNA hypoxia response elements (rHRE) in their 3’UTR. (B) Steady-state or “normal” DDX28 levels observed in normoxia contribute to the normoxic decrease in HIF-2α protein levels through binding with DDX28 and shifts the balance away from the formation of an activating complex with eIF4E2, repressing this non-canonical cap-dependent translation pathway. Since depleting DDX28 in hypoxia activates eIF4E2-dependent translation even further, we propose that some DDX28 is required in hypoxia to restrain this oncogenic pathway.

While depletion of DDX28 significantly increased the ability of eIF4E2 to bind m^7^GTP in both normoxia and hypoxia, it was surprising that this increase was greater in normoxia than in hypoxia (Fig. 2). This could indicate that DDX28 has increased relevance as a suppressor of eIF4E2 activity in normoxia, but while the polysome-associated eIF4E2 increased in DDX28 depleted normoxic cells, it was not statistically significant (Fig. 3F). Further, the polysome association of eIF4E2 mRNA targets *EGFR* and *IGF1R* did not increase in normoxic DDX28 depleted cells relative to control (Fig. 4B). In hypoxia, the effects of DDX28 depletion were not only observed on eIF4E2 m^7^GTP and polysome association, but here *EGFR* and *IGF1R* mRNAs were significantly more associated with polysomes relative to control (Fig. 4A). We speculate that depleting DDX28 in normoxia may increase the very low levels of HIF-2α to levels still undetectable via western blot, but enough to significantly increase the cap-binding potential of eIF4E2. The relative increase in HIF-2α in normoxic DDX28 depleted cells relative to control may be greater than that in hypoxic cells, but there could be an unmet requirement in normoxia for a threshold amount of total HIF-2α protein to efficiently activate eIF4E2 and the translation of its mRNA targets. Unexpectedly, the increase in nuclear HIF-2α upon DDX28 depletion in hypoxia did not produce subsequent increases to the transcription of its target genes (Fig. 5). We speculate that perhaps in hypoxic control cells, the HIF-2α DNA binding sites are saturated. Conversely, the cytoplasmic HIF-2α targets (i.e. eIF4E2, DDX28) are likely not saturated due to the much lower levels of HIF-2α in this compartment relative to the nucleus.

Hypoxia decreases total DDX28 protein levels, but our data suggest that the remaining DDX28 is important to restrain the HIF-2α/eIF4E2 translational axis. This brings into to question why a negative regulator of this pathway would be in place, given that the expression of rHRE-containing mRNAs contributes to hypoxic survival [25]. Dozens of eIF4E2 mRNA targets have been identified such as *EGFR*, *IGF1R*, *PDGFRA*, *CDH22* [16, 17, 22, 24], and these have all been characterized as oncogenes [24, 40, 41]. Therefore, while important for hypoxic adaptation, tight regulation of this pathway is likely required to prevent neoplastic transformation. Indeed, mining the data in The Pathology Atlas within The Human Protein Atlas, we found that DDX28 is present in most tissues, but lost in the majority of cancers (n ≥ 3 patients per cancer type; available from v18.proteinatlas.org and https://www.proteinatlas.org/ENSG00000182810-DDX28/pathology) [42]. Conversely, the presence of DDX28 is listed as a favorable prognostic marker in cases of renal cell carcinoma. More than 80% of all clear cell renal cell carcinomas (CCRCC), the most common form of renal cancer, contain inactivating mutations in the *VHL* gene that stabilize the HIF-α subunits in a hypoxia-independent manner [43]. The normoxic stabilization of HIF-2α and activation of its oncogenic pathways that occurs in CCRCC could be antagonized by the presence of DDX28. This study was performed in U87MG glioblastoma cells, but since DDX28 appears to be expressed ubiquitously, it could function similarly in other tissues. We provide mechanistic insight into the regulation of HIF-2α and eIF4E2-directed hypoxic translation, and support for DDX28 as a tumor suppressor and prognostic marker to deepen our understanding of cancer progression.

## METHODS

### Cell Culture

U87MG human glioblastoma cells (HTB-14) were obtained from the American Type Culture Collection and maintained as suggested. Normoxic cells were maintained at 37 °C in ambient O_2_ levels (21%) and 5% CO_2_ in a humidified incubator. Hypoxia was induced by culturing at 1% O_2_ and 5% CO_2_ at 37 °C for 24 h, unless otherwise stated, using an N_2-_balanced Whitley H35 HypOxystation.

### Generation of Stable Cell Lines

Two unique OmicsLink™ shRNA expression vectors (Genecopoeia) were used to target the coding sequence of human DDX28 [HSH014712-3-nU6 sequence 5’-ggtggactacatcttagag-3’, HSH014712-3-nU6 sequence 5’-acgctgcaagattacatcc-3’]. A non-targeting shRNA was used as a control. U87MG cells stably expressing C-terminal 3x FLAG tagged DDX28 were generated by transfecting cells with the OmicsLink™ pEZ-M14 EX-A3144-M14 expression vector encoding the human DDX28 coding sequence (Genecopoeia). Selection was initiated 48 h post-transfection using 1 μg/mL puromycin or 400 μg/mL G418, respectively, and single colonies were picked after seven days.

### Western Blot analysis

Standard western blot protocols were used. Primary antibodies: anti-eIF4E2 (Genetex, GTX82524), anti-DDX28 (Abcam, ab70821), anti-eIF4E (Cell Signaling, C46H6) anti-GAPDH (Cell Signaling, D16H11), anti-RPL5 (Abcam, ab137617), anti-HIF-2α (Novus, NB100-122), anti-FLAG (Sigma, F1804), anti-α-tubulin (GeneTex, GT114), anti-lamin a/c (Cell Signaling, 2032), anti-GFP (Abcam, ab290), anti-HA (Santa Cruz, Y-11), anti-β-Actin (Genetex, GT5512), and anti-EGFR (Proteintech, 18986-1-AP).

### Polysome Profiling and analysis

Performed as described previously [17]. The total eIF4E2 or eIF4E signal was quantified by densitometry using Bio-Rad Image Lab software, and the percentage of eIF4E2 or eIF4E present in monosome and polysome fractions relative to the total signal was calculated.

### RNA Isolation and quantitative RT-PCR

RNA was extracted from polysome fractions and qRT-PCR performed for Fig. 4A-B and E as previously described [17]. RNA extracted from cells for Fig. 4D and Fig. 5B using RiboZol™ as per manufacturer’s instructions. RNA (4 μg) was reverse transcribed using the high-capacity cDNA reverse transcription kit (Applied Biosystems). Primer sequences used (5’-3’): *CITED2*, CCT AAT GGG CGA GCA CAT ACA (forward) and CGT TCG TGG CAT TCA TGTT (reverse); *EEF2*, Forward TTC AAG TCA TTC TCC GAG A and Reverse AGA CAC GCT TCA CTG ATA; *EGFR*, GGA GAA CTG CCA GAA ACT GAC (forward) and GGG GTT CAC ATC CAT CTG (reverse); *EPO*, TGG AAG AGG ATG GAG GTC GG (forward) and AGA GTG GTG AGG CTG CGA A (reverse); *IGFBP3*, GCG CCA GGA AAT GCT AGT G (forward) and AAC TTG GGA TCA GAC ACC CG (reverse); *IGF1R*, CCA TTC TCA TGC CTT GGT CT (forward) and TGC AAG TTC TGG TTG TCG AG (reverse); *GAPDH*, GTC AAG GCT GAG AAC GGG A (forward) and CAA ATG AGC CCC AGC CTT C (reverse); *HSP90AB1*, TGT CCC TCA TCA TCA ATA CC (forward) and TCT TTA CCA CTG TCC AAC TT (reverse); *ITPR1*, CGG AGC AGG GTA TTG GAA CA (forward) and GGT CCA CTG AGG GCT GAA AC (reverse); *LOXL2*, CCC CCT GGA GAC TAC CTG TT (forward), GGA ACC ACC TAT GTG GCA GT (reverse); *OCT4*, GAT GTG GTC CGA GTG TGG TTC (forward) and TTG ATC GCT TGC CCT TCT G (reverse); *RPLP0*, AAC ATC TCC CCC TTC TCC (forward) and CCA GGA AGC GAG AAT GC (reverse); RPL13A, CAT AGG AAG CTG GGA GCA AG (forward) and GCC CTC CAA TCA GTC TTC TG (reverse). Relative fold change in expression was calculated using the ΔΔCT method, and transcript levels were normalized to *RPLP0* and either *RPL13A* or *GAPDH*.

### Immunoprecipitation and vectors

Exogenous expression vectors used: FLAG-GFP-HIF-2α in a pAdlox backbone was a gift from Dr. Stephen Lee (Miami), HA-HIF1alpha-pcDNA3 was a gift from William Kaelin (Addgene plasmid # 18949; http://n2t.net/addgene:18949; RRID:Addgene_18949), and FLAG-eIF4E2 was a gift from Dong-Er Zhang (Addgene plasmid # 17342; http://n2t.net/addgene:17342; RRID:Addgene_17342). Control vectors were of the same backbone and tag without a gene insert. Cells were transfected with 4 μg DNA complexed with 20 μg polyethylenamine (PEI) diluted in 600 μL of lactate buffered saline (20 mM sodium lactate and 150 mM NaCl, pH 4.0). The DNA/PEI complexes were diluted in 2.4 mL DMEM without FBS or antibiotics and added to cells at 37 °C for 8 h followed by replenishing with complete media. Cells were lysed after 48 h in 200 μL lysis buffer [10 mM Tris-HCl (pH 7.4), 150 mM NaCl, 0.5 mM EDTA, 0.5% NP-40, 1X protease inhibitor cocktail (New England Biolabs)]. Lysates were centrifuged at 12,000 rpm for 10 min at 4 °C and diluted with 500 μL of dilution buffer [10 mM Tris-HCl (pH 7.4), 150 mM NaCl, 0.5 mM EDTA, 1X protease inhibitor cocktail]. Protein (1 mg/mL) was incubated with 25 μL of GFP-Trap magnetic micro beads (ChromoTek), or anti-FLAG M2 magnetic beads (Sigma), or anti-HA magnetic beads (Pierce). Only GFP-Trap required pre-blocking with 3% BSA in TBS [50 mM Tris, 150 mM NaCl, pH 7.4] and washed as per the manufacturer’s instructions. Immunoprecipitation was carried out for 1 h at 4 °C with rotation. The GFP beads were washed four times with a more stringent wash buffer [10 mM Tris-HCl (pH 7.4), 500 mM NaCl, 0.5 mM EDTA, 0.1% NP-40] while FLAG and HA beads were washed four times with TBS. Proteins were eluted at 95 °C in 1X Laemmli sample buffer. Whole cell lysate (25 μg) was used as the input.

### Analysis of Cap-Binding Proteins

Performed as previously described [39]. The eIF4E2 signal in input and cap-elution lanes was quantified by densitometry using Bio-Rad Image Lab software. The eIF4E2 signal in the DDX28 knockdown cap-elution lane relative to control was normalized to the eIF4E2 input signal ratio.

### Cellular fractionation

After 24h of hypoxia, cells were lysed in 400 μL harvest buffer [10 mM HEPES, 50 mM NaCl, 500 mM sucrose, 0.1 mM EDTA, 10 mM DTT, 2 mM NaF, 0.5% Triton-X100, 1X protease inhibitor cocktail]. Lysates were centrifuged at 8000 rpm for 10 min at 4 °C to pellet nuclei. The supernatant was collected as the cytoplasmic fraction and the nuclear pellet was washed twice with 800 μL nuclear wash buffer [10 mM HEPES, 10 mM KCl, 0.1 mM EDTA, 0.1 mM EGTA, 10 mM DTT, 2 mM NaF, 1X protease inhibitor cocktail] centrifuging at 13,000 rpm at 4 °C for 5 and 10 min following washes. The nuclear pellet was resuspended in 400 μL RIPA buffer [20 mM Tris-HCl (pH 7.5), 10 mM NaCl, 1% NP-40, 0.1% SDS, 0.5% sodium deoxycholate, 1 mM EDTA, 10 mM DTT, 2 mM NaF, 1X protease inhibitor cocktail] and rotated at 4 °C for 15 min. Insoluble proteins were pelleted by centrifugation at 13,000 rpm for 10 min at 4 °C and the supernatant was collected as the nuclear fraction. Equal volumes of cytoplasmic and nuclear samples were mixed with 1X Laemmli sample buffer and boiled at 95 °C for 90 sec for western blot analysis.

### Viability assay

For each indicated time point, 10,000 cells per well were plated in triplicate in a 24-well plate. The following day (day 0), cells were incubated at their indicated oxygen concentrations, and following each 24 h increment, cells were washed once with PBS and stained with 400 μL of 1% crystal violet solution prepared in 20% methanol, with gentle rocking for 20 min at room temperature. Cells were gently washed with water to remove excess stain. Plates were air-dried overnight, and 400 μL of 10 % acetic acid was added to each well and incubated on a shaker for 20 min at room temperature for de-staining. The absorbance at 595 nm was measured using the ThermoMax microplate reader (Molecular Devices).

### Cell proliferation assay

Cells (250,000) were seeded on coverslips and incubated at their indicated oxygen concentrations for 24 h prior to treatment with 10 μmol/l bromodeoxyuridine (BrdU) cell proliferation labeling reagent (Sigma) for 1 h. Cells were washed with PBS and fixed in cold methanol for 10 min. Excess methanol was removed by washing for 5 min with PBS, and coverslips were incubated with 1:100 primary anti-bromodeoxyuridine antibody (RPN202; GE Healthcare) in the dark for 1 h. Coverslips were washed three times for 5 min with PBS and incubated with 2 μg/mL goat anti-mouse Alexa Fluor 555 secondary antibody (Invitrogen,), in the dark for 1 h at 37 °C. Cells were counterstained with Hoechst (1 μg/mL) for 5 min, and coverslips were mounted on microscope slides using ProLong Gold antifade reagent (Invitrogen). Cells were imaged with the Nikon eclipse Ti-S inverted microscope. An average of 200 cells were assessed for positive BrdU labeling per biological replicate.

### Statistical analyses

Results are expressed as means ± standard error of the mean (s.e.m) of at least three independent experiments. Experimental data were tested using unpaired two-tailed Student’s t-test when only two means were compared, or a one-way ANOVA followed by Tukey’s HSD test when three or more means were compared. P < 0.05 was considered statistically significant using GraphPad Prism.

## Abbreviations

BrdU: bromodeoxyuridine
CCRCC: clear cell renal cell carcinoma
DDX28: DEAD box protein 28
EEF2: Eukaryotic Translation Elongation Factor 2
EGFR: Epidermal Growth Factor Receptor
eIF4E: eukaryotic initiation factor 4E
HIF: Hypoxia-inducible factor HSP90ab1, Heat Shock Protein 90 Alpha Family Class B Member 1
IGF1R: Insulin Like Growth Factor 1 Receptor
PEI: polyethylenamine
rHRE: RNA Hypoxia Response Elements

## AUTHOR CONTRIBUTIONS

SLE, OB, and EJS performed experiments. SLE and JU planned experiments, analyzed data and wrote the manuscript. JU designed the study and provided all the resources.

## ACKNOWLEDGEMENTS

This work was funded by the Natural Sciences and Engineering Council of Canada (NSERC) grant number 04807 to J.U. S.L.E. was supported by the NSERC Canada Graduate Scholarships-Master’s (CGS M).

