## Supplemental Figure 1 for "DEAD-box protein family member DDX28 is a negative regulator of HIF-2α and eIF4E2-directed hypoxic translation"

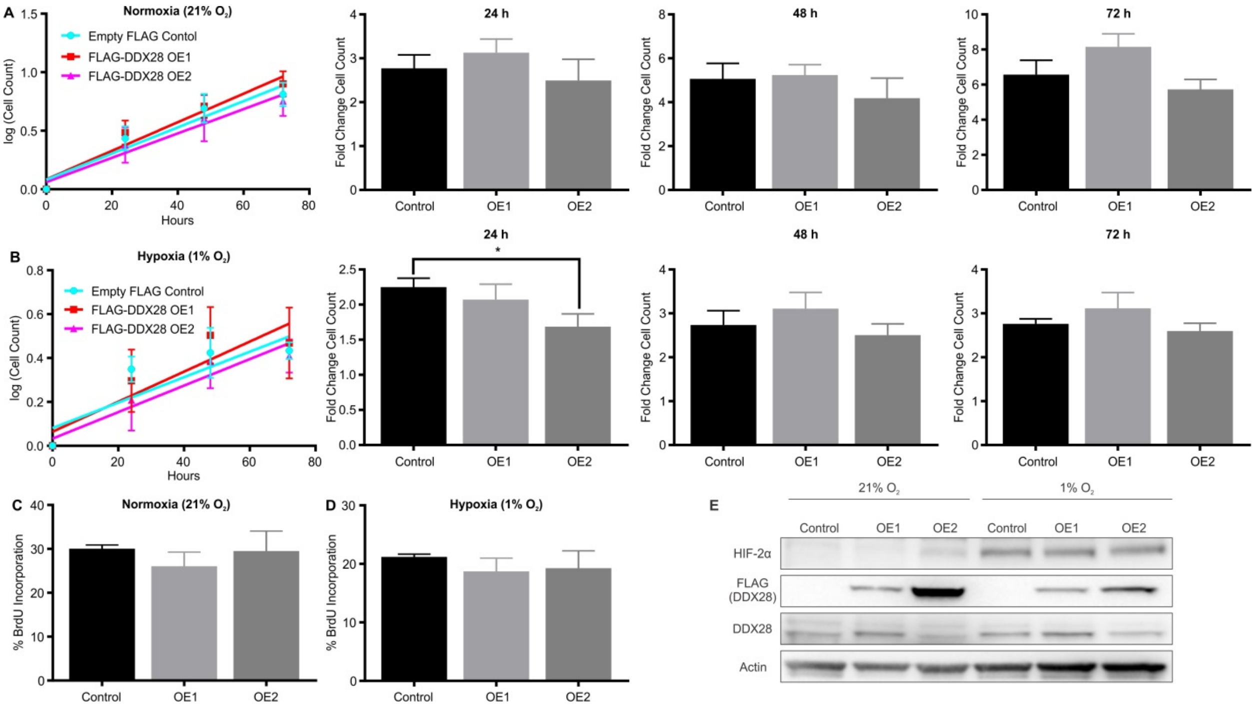


**Figure S1.** Overexpression of DDX28 does not alter cell viability and proliferation in normoxia and hypoxia. Viable cell counts were measured with crystal violet staining after 24 h, 48 h, and 72 h in 21% $O_2$ normoxia (A) and 1% $O_2$ hypoxia (B) for control cells stably expressing an empty FLAG vector or two clones stably overexpressing (OE) exogenous FLAG-DDX28 (OE1 and OE2). Proliferation was measured as % BrdU-positive control and DDX28 OE cells after 24 h in normoxia (C) or hypoxia (D). Data ($n \geq 3$), mean ± s.e.m. * represents $p < 0.05$ using an unpaired two-sample t-test. E, Western blot of HIF-2α, FLAG-DDX28, and endogenous DDX28 in normoxic and hypoxic control cells and DDX28 OE cells. Actin used as a loading control. Experiments performed in U87MG glioblastoma.
